# Testing the reliability of novel Voxel Placement approaches for Magnetic Resonance Spectroscopy

**DOI:** 10.64898/2026.08.11.744164

**Authors:** Harleen Chhabra, Melina Hehl, Koen Cuypers, Ulrike Dydak, Michael A. Nitsche, Erhan Genç, Michael Burke

## Abstract

**Background:** Single-voxel magnetic resonance spectroscopy (MRS) is a non-invasive method for measuring clinically and cognitively relevant metabolites. Reliable measurements require precise voxel placement across sessions and participants. We developed a scanner-console-based approach to improve voxel placement precision.

**Methods:** In a crossover design (n=7; six sessions each), we compared test–retest reliability of three voxel placement methods in a reference benchmark (left parietal cortex) and a technically challenging region (left ventromedial prefrontal cortex). Methods included (1) conventional anatomy-based placement, (2) mask-guided real-time positioning (MGRP), and (3) semiautomated session-locked voxel repositioning (SSVR). Resting-state MRS data were acquired using PRESS and MEGA-PRESS. Within-subject reliability of voxel placement and metabolite concentrations, namely, total N-acetylaspartate (tNAA), total Creatine (tCr), GABA (gamma-aminobutyric acid), and Glx (glutamate + glutamine) are reported using the coefficient of variation (CV), the intraclass correlation coefficient (ICC), minimal detectable change (MDC), and the spatial overlap.

**Results:** SSVR markedly improved voxel placement reliability, increasing spatial overlap (up to 88%) and achieving near-perfect geometric reproducibility (ICC = 0.99) compared to conventional anatomy-based placement and MGRP. SSVR improved tissue composition consistency and reduced metabolite variability in the technically challenging region (variability reduction of ∼70% tCr, ∼59% tNAA, and ∼51% Glx) while further refining already stable measurements in the benchmark region (tNAA from ∼15% to ∼10%).

**Conclusion:** Both MGRP and SSVR improved voxel placement and metabolite measurement reproducibility compared with conventional anatomy-based placement. SSVR further enhanced within-subject reproducibility across repeated sessions, particularly in the technically challenging region, providing a robust approach for longitudinal single-voxel MRS studies.

## 1. Introduction

Magnetic resonance spectroscopy (MRS) is a non-invasive technique that is used to quantify the metabolite and neurotransmitter concentration in the human brain, either during rest or functionally active brain states (Buonocore and Maddock, 2015; Mullins, 2018). MRS can measure metabolites like *N*-acetyl aspartate (NAA), creatine, choline, glutamine/glutamate (Glx), myo-inositol, lactate, and γ-aminobutyric acid (GABA), which are both cognitively and clinically relevant (Oz et al., 2014; Stanley and Raz, 2018). Unlike functional magnetic resonance imaging (fMRI), which acquires whole-brain blood oxygenation level-dependent (BOLD) data without requiring predefined regions, single-voxel spectroscopy (SVS) relies on prior anatomical or functional localization landmarks for data acquisition - an irreversible decision not correctable during post-processing. SVS is widely used for its high signal-to-noise ratio (SNR), which enables high-quality spectra. Conclusions drawn from MRS studies are usually based upon the functional role of a particular anatomical structure, and there is generally an implicit assumption that voxel placement is accurate and reproducible. Furthermore, the selection of a brain region for SVS is highly dependent on the research question, and therefore, it is crucial that the region of voxel placement is identified precisely and the placement is reliable with high accuracy across multiple sessions and study participants.

The most used voxel placement method is conventional anatomy-based placement (Bai et al., 2017; Dou et al., 2015), which depends heavily on operator expertise and is sensitive to inter-individual anatomical variability. One possible reason why this method remains widely used is that many MRI scanner console environments do not support prior mapping and marking of brain regions that could then guide voxel placement during the actual acquisition. These challenges are exacerbated with smaller volumes of interest, multi-session and multi-centre designs, and can affect associated biochemical measurements (Öz et al., 2020). While anatomy-based placement may be effective in regions with clear anatomical landmarks, like the hand knob region for the motor cortex (Yousry et al., 1997), this approach easily fails in regions with substantial functional and cytoarchitectural variability, like the prefrontal and parietal regions (Rajkowska and Goldman-Rakic, 1995). Additionally, anatomical variability contributes to poor voxel overlap in test–retest scenarios (Bai et al., 2017; Lee et al., 2013). Moreover, even when spatial consistency is achieved using visible landmarks, it does not ensure sampling of functionally equivalent regions across individuals, as underlying functional and cytoarchitectural differences are not directly observable.

Although automated voxel placement methods have been previously attempted, like using the scanner Auto-Align function and custom-built Linux-based Automated Voxel Placement (AVP) (Dou et al., 2015; Woodcock et al., 2018), which improved reproducibility compared to manual anatomy-based placement approaches, they remain limited by restricted validation across brain regions, reliance on template-based or center-of-gravity (CoG) approaches for registration, and residual sensitivity to inter-individual anatomical variability (Bishop et al., 2023; Dou et al., 2015; Hancu et al., 2005; Storrs, 2010; Woodcock et al., 2018). Furthermore, even with improved spatial consistency, partial volume effects continue to introduce variability in tissue composition and metabolite estimates.

To address these limitations, we established a novel voxel placement approach and conducted this study comparing MRS reliability and reproducibility of voxel placement and metabolite concentrations within and between healthy participants at a reference benchmark region and a technically challenging region. The left parietal cortex was selected as the reference benchmark region, and the left ventromedial prefrontal cortex (vmPFC) as the technically challenging region. The vmPFC was chosen as a technically challenging region due to its proximity to sinus cavities, where susceptibility artifacts, noise from the frontal bone, and cerebrospinal fluid degrade MRS signal quality. Additionally, prior vmPFC studies have relied only on anatomical landmarks (e.g., genu of the corpus callosum, anterior cingulate cortex) with varying voxel sizes, leading to inconsistent placement across studies (Chen et al., 2017; Zhang et al., 2016).

We established and compared two new voxel placement pipelines with the traditional anatomical-landmark placement method: 1) the mask-guided real-time positioning (MGRP) done for real-time visual guidance on the MRI scanner console using a subject-specific mask (anatomical parcellation-/regional activation-based mask), and 2) the semiautomated session-locked voxel repositioning (SSVR) done for automated registration and re-localization of the prior-session voxel mask that could significantly enhance the reliability and reproducibility of voxel placement across sessions. Both methods utilize T1-weighted images with region- or session-specific masks, rather than relying solely on anatomical landmarks, CoG, or template approaches. While these placements are effective in regions like the dorsolateral prefrontal and cingulate cortex due to their anatomy (Bishop et al., 2023; Dou et al., 2015; Hancu et al., 2005; Storrs, 2010; Woodcock et al., 2018), it is less suitable for our chosen technically challenging region, where susceptibility artifacts can distort signal localization. In contrast, mask-based approaches (MGRP, SSVR) enable weighted voxel placement that prioritizes metabolically relevant tissue and maximizes gray matter overlap, potentially improving measurement consistency by accounting for subject-specific anatomical differences.

Furthermore, for both the reference benchmark and technically challenging region, one voxel each was placed the traditional anatomical-landmark and MGRP method. The voxel mask used for MGRP was derived from individual brain-parcellation. Additionally, for the technically challenging region another voxel was placed with the mask derived from task-based functional activation.

Reproducibility and reliability were assessed using the coefficient of variation (CV), intraclass correlation coefficient (ICC), minimal detectable change (MDC), and voxel spatial overlap. CV reflects relative variability and measurement stability [4]. ICC evaluates the ability to distinguish between individuals despite measurement variability but is limited when between-subject variability is low [3](Eftekhari et al., 2025). MDC represents the smallest detectable change beyond measurement error (typically at 95% confidence) (Pratt et al., 2025). Spatial overlap (global overlap) assesses the consistency of voxel placement across sessions and participants.

We hypothesized that the newly developed MGRP would improve voxel placement reliability relative to conventional anatomy-based placement in both regions, and that SSVR, in addition, would outperform both conventional anatomy-based placement and MGRP in a multi-session context. Similar improvements were expected for metabolite concentration reliability (tNAA, tCr, Glx, GABA).

## 2. Methodology

### 2.1 Participants

Seven (4 females, age = 30.43±3.99) healthy, non-smoking right-handed participants were recruited for this study. The participants were excluded if they were diagnosed with any neurological or psychiatric disease or had a history of such diseases, had a pacemaker or metal implants anywhere in the body, suffered from circulatory disturbance of the brain or cerebral haemorrhage, had evidence of epileptic seizures or the presence of epilepsy, had a history of head injury with injury to the skull bone or brain damage, alcohol, or drug dependence, and were either pregnant or lactating. A trained medical doctor assessed the inclusion and exclusion criteria that were self-reported by the participants. The protocol was approved by the ethics committee of the Leibniz Research Centre for Working Environment and Human Factors (No 117-2) and was in accordance with the Declaration of Helsinki (2013). All participants gave written consent for the study.

### 2.2 Study design and procedure

The study was conducted to compare the test-retest reliability of three voxel placement methods across the reference benchmark region and the technically challenging region. The left parietal cortex was used as the reference benchmark region, and left ventromedial prefrontal cortex was used as the technically challenging region (Fig. 1). The three voxel placement methods used in the study were: 1) the conventional anatomy-based placement done using the manual placement via anatomical landmarks, 2) the mask-guided real-time positioning (MGRP) done for real-time visual guidance using a subject-specific mask (anatomical parcellation-/regional activation-based mask), and 3) the semiautomated session-locked voxel repositioning (SSVR) done for automated registration and re-localization of the prior-session voxel mask (Fig. 2).

**Fig. 1:**
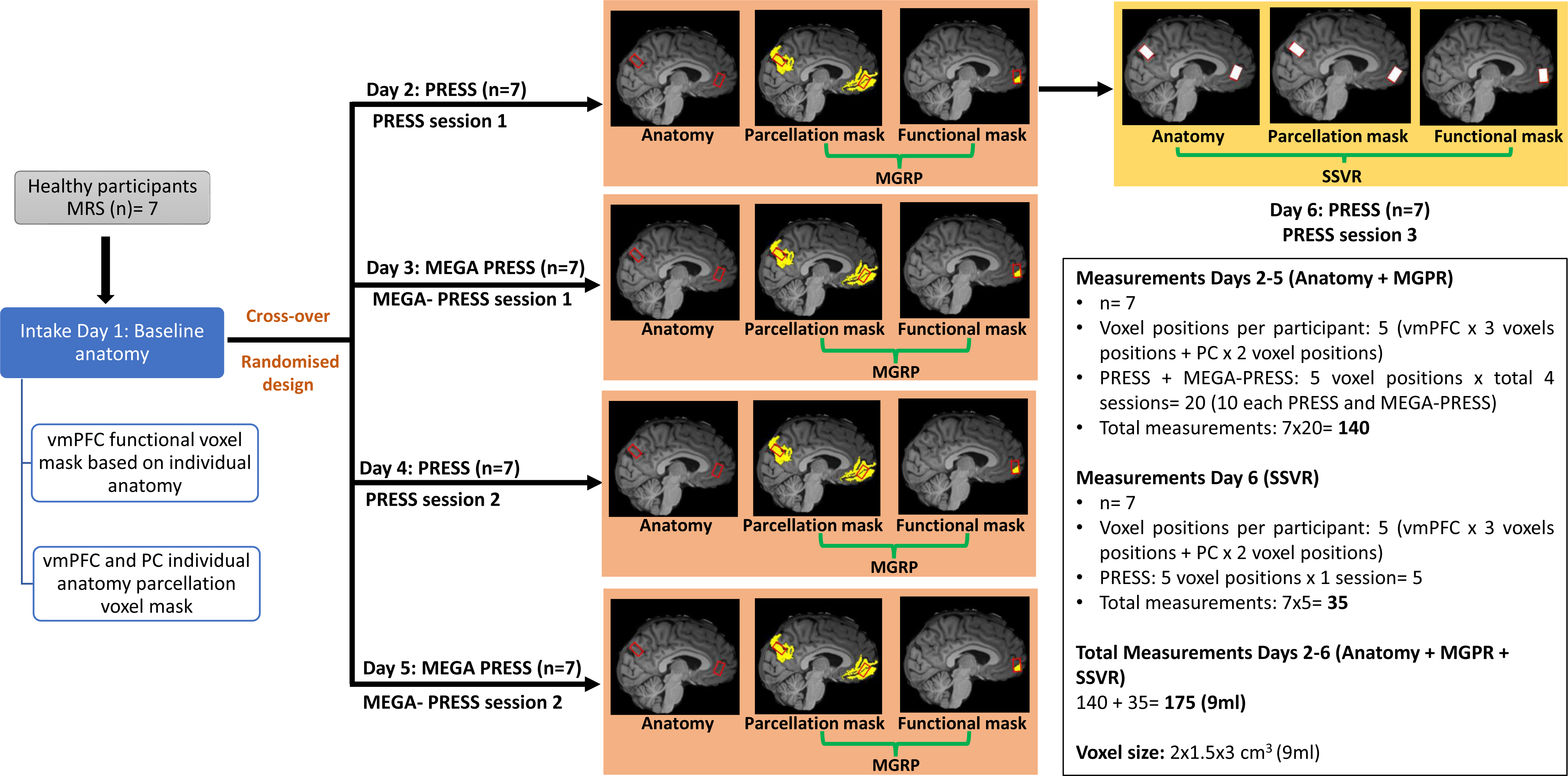
Study methodology. The six-day study design consisted of one intake session (day 1) followed by three days of PRESS MRS (days 2, 4, 6) and two days of MEGA-PRESS (days 3,5). MRS was acquired from the reference benchmark region (left PC: parietal cortex) and the technically challenging region (left vmPFC: ventromedial prefrontal cortex) with a voxel size of 2×1.5×3cm^3^ (see Fig S1). For the PC, the voxels were placed based on anatomy and individual parcellation mask. For the vmPFC, the voxels were placed based on anatomy, individual parcellation mask, and functional mask obtained from the Fullana et al. (2018) fear extinction meta-analysis (Fullana et al., 2018). For the conventional anatomy-based placement, the voxel was placed on the anatomical landmarks (Mahone et al., 2018; Silveira de Souza et al., 2011; Yang et al., 2015). For the mask-guided real-time positioning (MGRP), the individual anatomy parcellation was done using the Human Connectome Cortex Multimodal Parcellation (HCCMP) (Glasser et al., 2016) in FreeSurfer, and the functional activation mask was registered to the individual space. For the last PRESS MRS session (day 6), the voxel was placed using the in-house semiautomated session-locked voxel repositioning (SSVR) method at both the PC and vmPFC. This methodology generated a total of 175 measurements across all seven participants.

**Fig. 2:**
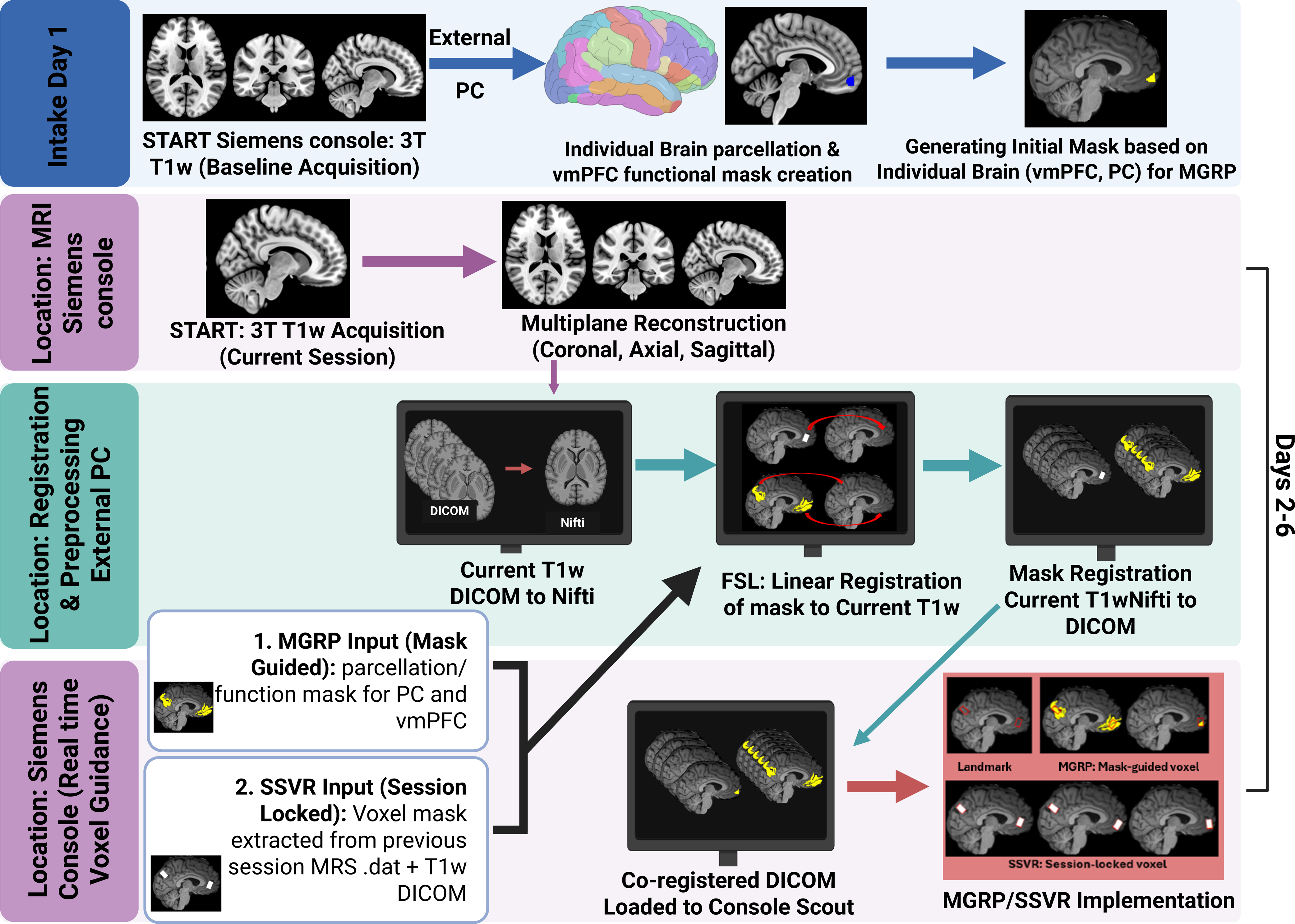
Mask-guided real-time positioning (MGRP) and Semiautomated session-locked voxel repositioning (SSVR) workflow. MRS data were acquired from the reference benchmark region (left parietal cortex: PC) and the technically challenging region (left ventromedial prefrontal cortex: vmPFC) using a voxel size of 2×1.5×3 cm³ (see Fig S1). In the MGRP pipeline, session T1-weighted (T1w) DICOM images were transferred to the registration and preprocessing computer, converted to NIfTI, and linearly co-registered to the baseline T1w images using FSL. Anatomical parcellation and/or fMRI activation masks were registered to the session T1w image, converted back to DICOM, and used on the scanner console for real-time voxel placement guidance. In the SSVR pipeline, subject-specific voxel masks were generated from the first MRS session by extracting voxel geometry from T1w images and spectroscopy data (.dat). These masks were registered to subsequent session T1w images using the same processing steps as MGRP and used to guide consistent, session-locked voxel placement across both regions. (Created in https://BioRender.com)

The study used a crossover design with six imaging sessions per participant (see Fig.1, Fig.S1). Structural MRI data were acquired during the intake session (Day1). Resting-state PRESS data were collected in days 2, 4, and 6, and resting-state MEGA-PRESS was collected in days 3 and 5. For each participant, the MRS sessions were conducted within 3 weeks. Two separate sequences were implemented because extracting glutamate from long-TE MEGA-PRESS sequences leads to systematic underestimation and higher measurement error due to T_2_ signal decay, making it unsuitable as a reliable benchmark (Cheng et al., 2021).Consequently, separate short-TE PRESS scans were performed to ensure high precision for glutamate and to mitigate the regional noise that already challenges GABA quantification within the MEGA-PRESS acquisition (Duda et al., 2021; Wang et al., 2024). The ^1^H MRS data were acquired from the reference benchmark region and the technically challenging region (see section 2.2, for details on voxel placement definitions) with a voxel size of 2×1.5×3cm^3^ (= 9ml). In addition, structural MRI was repeated at each session to enable real-time voxel planning and data collection. All imaging data were acquired at IfADo in Dortmund, Germany, using a 3T Siemens Prisma scanner with a 64-channel head coil (ve syngo MR E11) (Fig. 1). The study was designed with two primary experimental objectives: (i) to evaluate and compare the reliability of voxel placement within and between participants across the three voxel placement methods, and (ii) to assess the test-retest reliability of tNAA, tCr, GABA+, and Glx within and between participants’ resting-state MRS sessions.

### 2.3 Structural MRI (sMRI)

For co-registration of functional data, brain parcellation, and planning of MRS voxel placement in this study, T1-weighted sagittal anatomical images were acquired (TR = 2530 ms, TE = 2.36 ms, flip angle = 7°, 176 slices, matrix size = 256 mm x 256 mm, resolution = 1×1×1 mm^3^, duration ∼6 min). Images were automatically placed based on a Siemens-built-in atlas (AutoAlign) (ve syngo MR E11) and were aligned to the AC-PC line. This atlas-based placement of imaging planes allowed for maximized similarity of images used for voxel placement between sessions.

### 2.4 Individual anatomy brain parcellation

For reconstructing the cortical surfaces of T1-weighted images for all individuals, surface-based methods in FreeSurfer (http://surfer.nmr.mgh.harvard.edu, version 20 6.0.0) (Fischl, 2012) were used. The automated reconstruction steps included skull stripping, white and grey matter segmentation as well as reconstruction and inflation of the cortical surface. The brain parcellation was performed to define regions by the Human Connectome Project’s multi-modal parcellation (HCPMMP) (Glasser et al., 2016).

### 2.5 Voxel definitions for the reference benchmark region: left parietal cortex and the technically challenging region: left ventromedial prefrontal cortex (vmPFC) (See Fig. S1)

a. **Parietal conventional anatomy-based voxel:** The voxel was positioned laterally over the left parietal cortex. In the sagittal plane, the voxel was oriented parallel to the central sulcus, located posterior to the postcentral sulcus, and positioned with its posterior/inferior boundary just above the posterior cingulate cortex. In the axial plane, the voxel was centered over the left parietal lobe, lateral to the midline and posterior to the postcentral gyrus. In the coronal plane, the voxel was placed within the lateral parietal cortex while remaining superior to the posterior cingulate cortex(Silveira de Souza et al., 2011).
b. **Parietal individual anatomy brain parcellation-mask voxel:** T1-weighted images were processed with FreeSurfer using the HCPMMP atlas to obtain subject-specific parcellation. The parietal cortex region, defined in surface space and converted to volumetric space, included regions of interest (ROIs) 7Pm, 7PL, 7m, POS2, and 31pd (Glasser et al., 2016).
c. **vmPFC conventional anatomy-based voxel:** In the sagittal plane, the voxel was oriented parallel to the brainstem, with its centre aligned to the genu of the corpus callosum and positioned within the medial frontal region, while avoiding extension into the frontal bone and CSF. In the axial plane, the voxel was centred in the left vmPFC, while avoiding the orbital cavities (eye sockets) and minimizing contamination from surrounding air–tissue interfaces. In the coronal plane, the voxel was positioned with its dorsal boundary located anterior to the genu of the corpus callosum (Mahone et al., 2018; Yang et al., 2015).
d. **vmPFC individual anatomy brain parcellation-mask voxel:** Using FreeSurfer and the HCPMMP atlas, subject-specific vmPFC parcellations were generated and converted to volumetric space. The vmPFC region comprised ROIs s32, p32, 10v, 10r, a24, and 25 (Glasser et al., 2016).
e. **vmPFC fMRI functional activation-mask voxel:** A group-level vmPFC activation mask from a fear extinction paradigm in MNI space (Fullana et al., 2018) was warped to each participant’s structural T1-weighted image using FSL FNIRT (FSL version 6.0, RRID:SCR_002823), yielding subject-specific activation masks.

### 2.6 Mask-guided real-time positioning (MGRP): Real-time visual guidance using a subject-specific mask (anatomical parcellation-/regional activation-based mask)

During each scanning session, high-resolution T1-weighted (T1w) anatomical images were acquired in DICOM format and reconstructed across all three orthogonal planes on the Siemens MRI acquisition console. Images were sent from the Siemens console to an external PC, converted from DICOM to NIfTI, and co-registered to each participant’s baseline T1w scan using FSL to ensure alignment with prior anatomical and functional masks. Individualized masks were derived from structural parcellation (see 2.3, method b and d) and task-based fMRI activation maps (see 2.3, method e) (Fullana et al., 2018). For the reference benchmark region, a single parcellation mask was used, while for the technically challenging region, two masks were applied, one for parcellation and one for functional activation. These masks were registered to the session space, overlaid on T1w images, converted back to DICOM, and sent back to the Siemens scanner console to be used as real-time guides for voxel placement (Fig. 2). This pipeline ensured reproducible voxel positioning across sessions while maintaining sensitivity to individual anatomical differences and functional activation patterns.

### 2.7 Session-locked voxel repositioning (SSVR) for automated registration and re-localization of a prior-session voxel mask

Extending this MGRP approach, session-locked voxel masks were generated from the first MRS session by reconstructing voxel geometry from spectroscopy data and aligning voxel geometry with T1w DICOM images. For the subsequent session, these masks were registered to the current session’s T1w images using the same pipeline as used for the MGRP method (Fig. 2). This extended workflow allowed reproducible, subject-specific session-locked voxel positioning across subjects while preserving alignment with prior placements.

### 2.8 MRS Protocol

Data from vmPFC and PC were acquired. For voxel placement, structural T1-weighted images were acquired using a MPRAGE pulse sequence as described above in section 2.2. These images were then transferred to the external voxel planning PC using the DICOM store Service Class User (SCU) mechanism as implemented on the scanner. The T1-weighted images were then further processed to automatically identify and mark the vmPFC and parietal cortex (see below for details). The labelled images were then transferred to the MRI scanner console to guide voxel placement at the previously identified locations. The MRS shimming was performed using the FASTESTMAP (Fast, Automatic Shim Technique using Echo-planar Signal readout, Mapping Along Projections) (Gruetter and Tkác, 2000) approach. Both PRESS and MEGA-PRESS (C2P package) were acquired using the Center for Magnetic Resonance Research (CMRR) spectroscopy package sequences. For Glx, the Center for Magnetic Resonance Research (CMRR) spectroscopy package PRESS sequence (TR=2000ms, TE=30ms, 64 averages, duration ∼8min) was used. Water-suppressed and non-water-suppressed spectra (16 averages) was acquired for postprocessing from both voxel locations. For GABA, GABA-edited MEGA-PRESS (Mescher et al., 1998) as implemented in the CMRR’s C2P spectroscopy package was used (TR=2000ms, TE=68ms, 96 averages, 1.9ppm ON, 7.5ppm OFF, pulse width=58Hz, duration ∼11min). Water-suppressed spectra and non-water-suppressed spectra (16 averages) for postprocessing were acquired from both locations (See MRSinMRS Checklist (Lin et al., 2021) in the Supplementary document).

### 2.7 MRS data analysis

Spatial overlap of MRS voxels within-subjectacross multiple sessions, as well as across the three voxel placement methods, was quantified using similarity index/spatial overlap coefficients (Dice, 1945) and Euclidean distances (Gower, 1984). Comparable analyses were also performed to evaluate between-participant differences in voxel placement and spatial overlap across the different placement approaches. Resting-state MRS was analyzed using Osprey (Oeltzschner et al., 2020) for PRESS data to obtain Glx, tCr aand tNAA, and Gannet (Edden et al., 2014) for MEGA-PRESS data to obtain GABA+ (GABA + macromolecules) quantification.

- Euclidean distance from center (0,0,0) set to the Anterior Commissure (AC) of the brain: 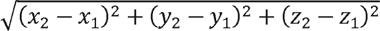 where x1, y1, z 1 is 0,0,0 and x2, y2, z2 is voxel centroid
- Global voxels overlap across individual participant sessions in subject space: 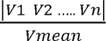 (V= voxel volume)

#### 2.7.1 PRESS analysis

Data were preprocessed and quantified using Osprey (v2.8.1), an open-source MRS toolbox running on MATLAB (R2021b) with an integrated LCModel fitting algorithm (Oeltzschner et al., 2020). The single-shot processing pipeline included coil combination, frequency and phase correction, eddy-current correction (via water reference), averaging, residual water removal, and spline-based baseline correction (knot = 0.5ppm). Spectral registration was done using the Robust spectral processing option, with spectral alignment done using L1Norm. Tissue segmentation of the gray matter (GM), white matter (WM), and cerebrospinal fluid (CSF) was performed using SPM12 within Osprey. Data quality metrics included tCr SNR, Cramer-Rao lower bounds (CRLB), and water linewidth (Supplementary Tables S4 and S6). SNR was calculated as the tCr peak height divided by noise standard deviation (−2 to 0 ppm), favouring robustness over peak integrals. Linewidth was defined as the full width at half maximum (FWHM) of the water peak (4.4–5.0 ppm).

#### 2.7.2 MEGA-PRESS analysis

Data were preprocessed and quantified using Gannet v3.3.2 (Edden et al., 2014). The preprocessing steps included zero-filling, eddy-current correction, and the application of a 3 Hz exponential line-broadening filter. Difference spectra were fitted within the 2.79 to 4.10 ppm range to quantify the GABA+ (3.0 ppm) and Glx (3.75 ppm) peaks using a three-Gaussian function with least-squares fitting. Metabolite concentrations were referenced to the water signal using a Gaussian–Lorentzian model between 3.8 and 5.6 ppm on the water-unsuppressed spectrum. Tissue segmentation (GM, WM, CSF) was performed using SPM12 within Gannet. Data quality metrics included were GABA+ SNR, fit error, and GABA+ FWHM (Supplementary Tables S3 and S5).

#### 2.7.3 Reliability analysis

Standard data quality metrics, including SNR, FWHM, LCModel Cramér–Rao lower bounds (CRLB) for PRESS, and Gannet MEGA-PRESS derived GABA+ fit error (FitErr), were used to reject the low-quality data. Data with SNR and/or FWHM ≥±2SD and CRLB and FitErr > 20% were removed from metabolite quantification. Quantified metabolites used for further analysis were tNAA, tCr, Glx from PRESS, and GABA+ from MEGA-PRESS. tNAA and tCr have high concentrations and are reported to be stable across sessions and individuals under healthy resting-state conditions (Caramanos et al., 2005). Whereas Glx and GABA+ have relatively low concentrations and are significantly influenced by brain state and noise levels in the data. Reproducibility and reliability of all parameters were quantified using the ICC, CV% and MDC. CV% and MDC were calculated for each subject and then meaned to give within-subject values.

Tests including the Pearson correlation coefficient, paired t tests, and Bland-Altman plots were not used in this study to test reliability. Paired t-tests and Bland-Altman plots only inform about the data agreement between repeated measurements at the group and individual levels, respectively, but do not quantify reliability. Likewise, the Pearson correlation coefficient only measures the strength of the linear association between repeated measurements but does not assess agreement between repeated measurements at the group and individual levels. Hence, these methods are not ideal for reliability analysis, which needs combined assessment of both the degree of correlation and agreement between measurements. In contrast, the intraclass correlation coefficient (ICC) provides a comprehensive measure of reliability by simultaneously accounting for both the consistency of measurements across individuals and the agreement between repeated measurements within individuals, CV provides a complementary measure of within-subject measurement variability, and MDC provides a measure of absolute reliability by estimating the smallest change that exceeds the measurement error at the individual level. ICC was estimated using SPSS version 25 (SPSS Inc, Chicago, IL) and CV% and MDC were calculated with RStudio 2026.07 (Team, 2020).

- Formulas used
- ICC estimates and their 95% confident intervals were calculated based on single rater mean-measurements, absolute-agreement, 2-way mixed-effects model.
- Within-subjects Coefficient of variance (CV%): 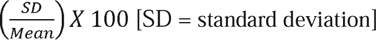
- Within-subjects Minimal detectable change (MDC) = 1.96 x SEM x √2 [SEM: Standard error of means]

## 3 Results

### 3.2 Voxel placement: Global Overlap and Euclidean Distance at the reference benchmark region and the technically challenging region using conventional anatomy-based placement, MGRP, and SSVR methods

At the reference benchmark region, the mean voxels overlap within and between subjects was 0.64 (value of 1 indicating the maximum overlap) for both the conventional anatomy-based placement and MGRP methods. The Euclidean distance from the centre had a CV% of 6.10 and 5.31, and ICC of 0.89 and 0.72 for the anatomical landmark and MGRP methods, respectively (see Fig. 3a, 3b and Fig. 4a, 4c global overlap in individual brains and mean within-subject values of voxel global overlap and Euclidean distance respectively). The SSVR method significantly improved the mean global voxel overlap within subjects to 0.85 for both conventional anatomy-based placement (anatomical voxel compared to SSVR anatomical voxel) and MGRP (MGRP voxel compared to SSVR MGRP voxel) methods. Furthermore, the SSVR method improved the Euclidean distance ICC to 0.99, while the CV% reduced for both the anatomical landmarks (1.16) and a parcellation mask voxel (1.21) (See Table 1, Fig. 3c and and Fig. 4a, 4c).

**Fig. 3:**
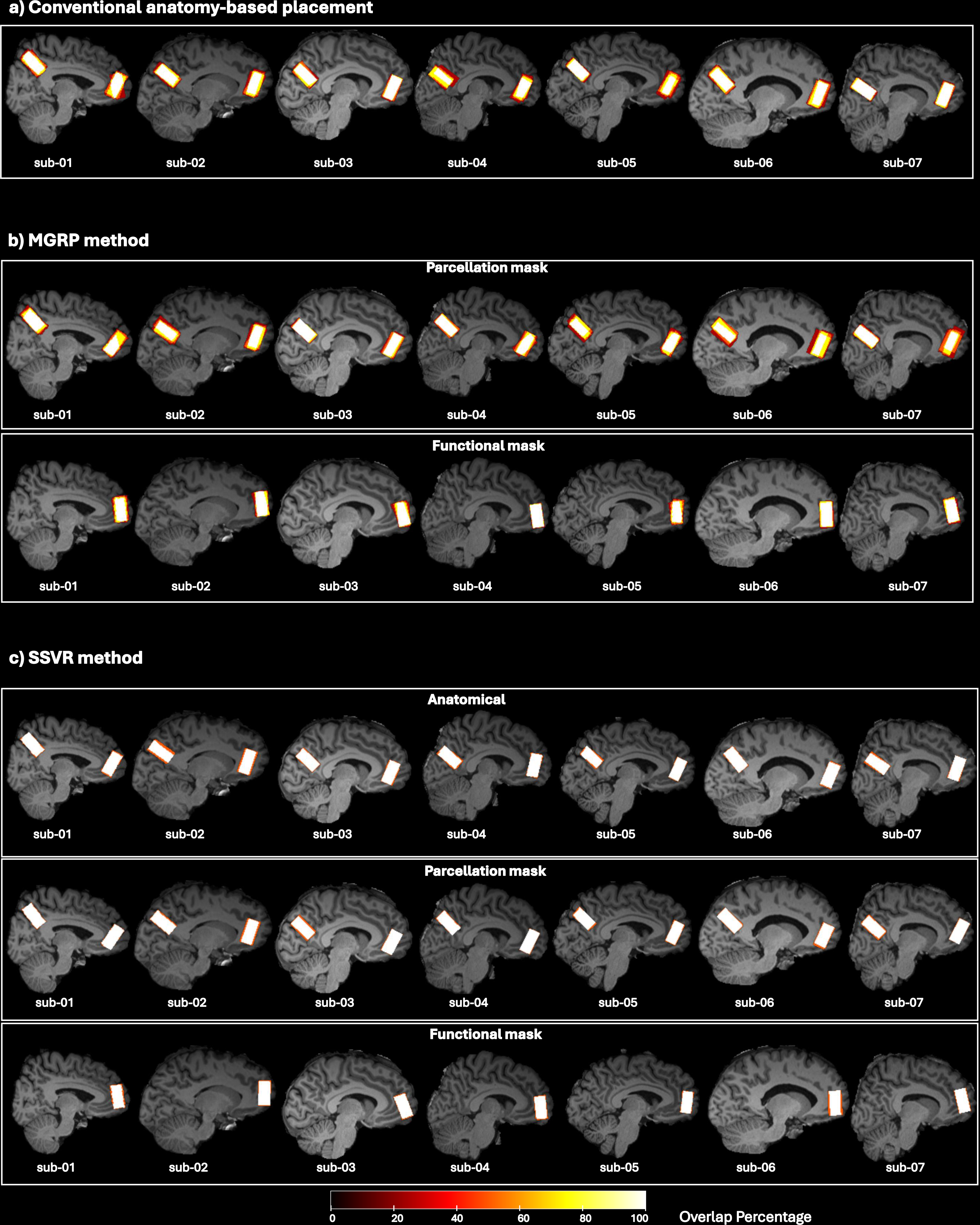
Group voxels global overlap (n=7) overlaid on the individual brain. Figure shows the global overlap for the MRS voxels placed at the reference benchmark region (left parietal cortex) and the technically challenging region (left ventromedial prefrontal cortex) using a) the conventional anatomy-based placement, b) Mask-guided real-time positioning (MGRP) methods and, c) the Semiautomated Session-locked voxel repositioning (SSVR). The color bar indicates the overlap percentage, with brighter colors indicating larger overlap.

**Fig. 4:**
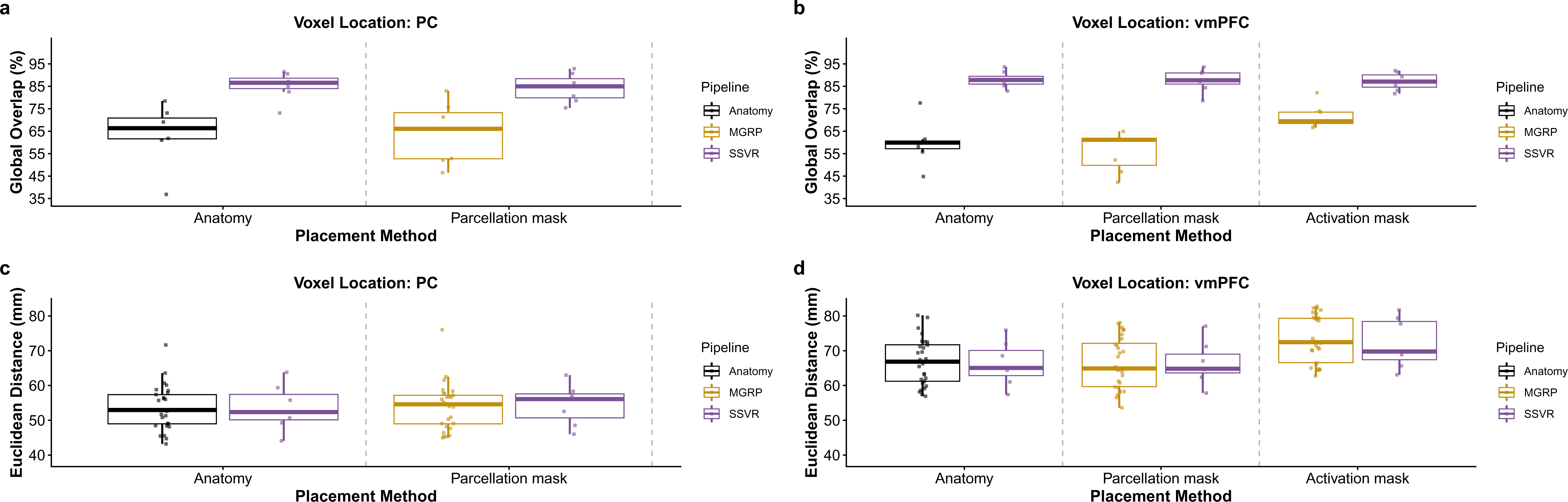
Voxel mean global overlap and mean euclidean distance (n=7). Figures a) and c) show the global overlap and Euclidean distance, respectively, for the MRS voxels placed at the reference benchmark regions (left PC) using the conventional anatomy-based placement, Mask-guided real-time positioning (MGRP), and Semiautomated Session-locked voxel repositioning (SSVR) methods. Figures b) and d) show the global overlap and Euclidean distance, respectively, for the MRS voxels placed at the technically challenging region (left vmPFC), also using the conventional anatomy-based placement, MGRP, and SSVR methods.

**Table 1:** Voxel global overlap and test-retest reliability for voxel tissue segmentation and voxel Euclidean distance from centre as measured with interclass correlations (ICC) and coefficient of variance percentage (CV%) for the reference benchmark region (left parietal cortex) and the technically challenging region (left vmPFC: ventromedial prefrontal cortex). fGM: fraction grey matter, fWM: fraction white matter, fCSF: fraction cerebrospinal fluid, ED: Euclidean distance, MGRP: Mask-guided real-time positioning, SSVR: Semiautomated Session-locked voxel repositioning.

| Reference Benchmark region (left PC) |  |  |  |  |  |  |  |  |  |
| --- | --- | --- | --- | --- | --- | --- | --- | --- | --- |
| Voxel Placement<br>(n=7) | Global Overlap<br>(subject space) | fGM |  | fWM |  | fCSF |  | ED |  |
|  | Mean±SD | ICC | CV% | ICC | CV% | ICC | CV% | ICC | CV% |
| Anatomical | 0.64±0.13 | 0.84 | 4.69 | 0.81 | 9.69 | 0.88 | 20.08 | 0.89 | 6.10 |
| MGRP |  |  |  |  |  |  |  |  |  |
| Parcellation-mask | 0.64±0.14 | 0.85 | 4.23 | 0.77 | 10.43 | 0.92 | 16.91 | 0.72 | 5.31 |
| SSVR |  |  |  |  |  |  |  |  |  |
| Anatomical | 0.85±0.06 | 0.89 | 5.40 | 0.91 | 7.58 | 0.97 | 11.42 | 0.99 | 1.16 |
| Parcellation-mask | 0.84±0.06 | 0.91 | 2.45 | 0.87 | 6.00 | 0.91 | 20.82 | 0.99 | 1.21 |
| Technically challenging region (left vmPFC) |  |  |  |  |  |  |  |  |  |
| Anatomical | 0.60±0.10 | 0.78 | 2.72 | 0.74 | 7.59 | 0.60 | 32.27 | 0.87 | 4.42 |
| MGRP |  |  |  |  |  |  |  |  |  |
| Parcellation-mask | 0.56±0.09 | 0.67 | 3.78 | 0.67 | 7.23 | 0.51 | 22.22 | 0.74 | 4.26 |
| Activation-mask | 0.72±0.05 | 0.89 | 3.48 | 0.90 | 7.31 | 0.56 | 18.92 | 0.98 | 3.90 |
| SSVR |  |  |  |  |  |  |  |  |  |
| Anatomical | 0.88±0.04 | 0.92 | 1.20 | 0.93 | 3.92 | 0.94 | 14.45 | 0.99 | 2.07 |
| Parcellation-mask | 0.88±0.05 | 0.96 | 1.81 | 0.93 | 4.43 | 0.67 | 19.03 | 0.99 | 1.68 |
| Activation-mask | 0.87±0.04 | 0.98 | 1.36 | 0.85 | 5.24 | 0.74 | 14.94 | 0.99 | 0.72 |

At the technically challenging region, the mean within-subject voxel overlap varied for the conventional anatomy-based placement (mean=0.60), parcellation mask (mean=0.56), and activation mask (mean=0.72) placed using the MGRP method. The Euclidean distance from the centre had a CV% of 4.42, 4.26, and 3.90, and an ICC of 0.87, 0.74, and 0.98 at the anatomical landmarks, parcellation mask, and activation mask voxel, respectively, for the conventional anatomy-based placement and MGRP methods (see Fig. 3a, 3b and Fig. 4b, 4d global overlap in individual brains and mean within-subject values of voxel global overlap and Euclidean distance respectively). The SSVR method significantly improved the mean global voxel overlap within subjects, between 0.87 and 0.88 for all voxels. Furthermore, the SSVR method improved the Euclidean distance ICC to 0.99 and the CV% for the anatomical landmarks (2.07), parcellation mask (1.68), and activation mask (0.72) (See Table 1, Fig. 3c. Fig. 4b, 4d).

### 3.3 Voxel placement : Brain segmentation (fGM, fWM, fCSF)

At the reference benchmark region, the ICC and CV% values remain similar for the conventional anatomy-based placement, MGRP, and SSVR (refer to Tables 1 and S1, and Fig. 5a and 5b).

**Fig. 5:**
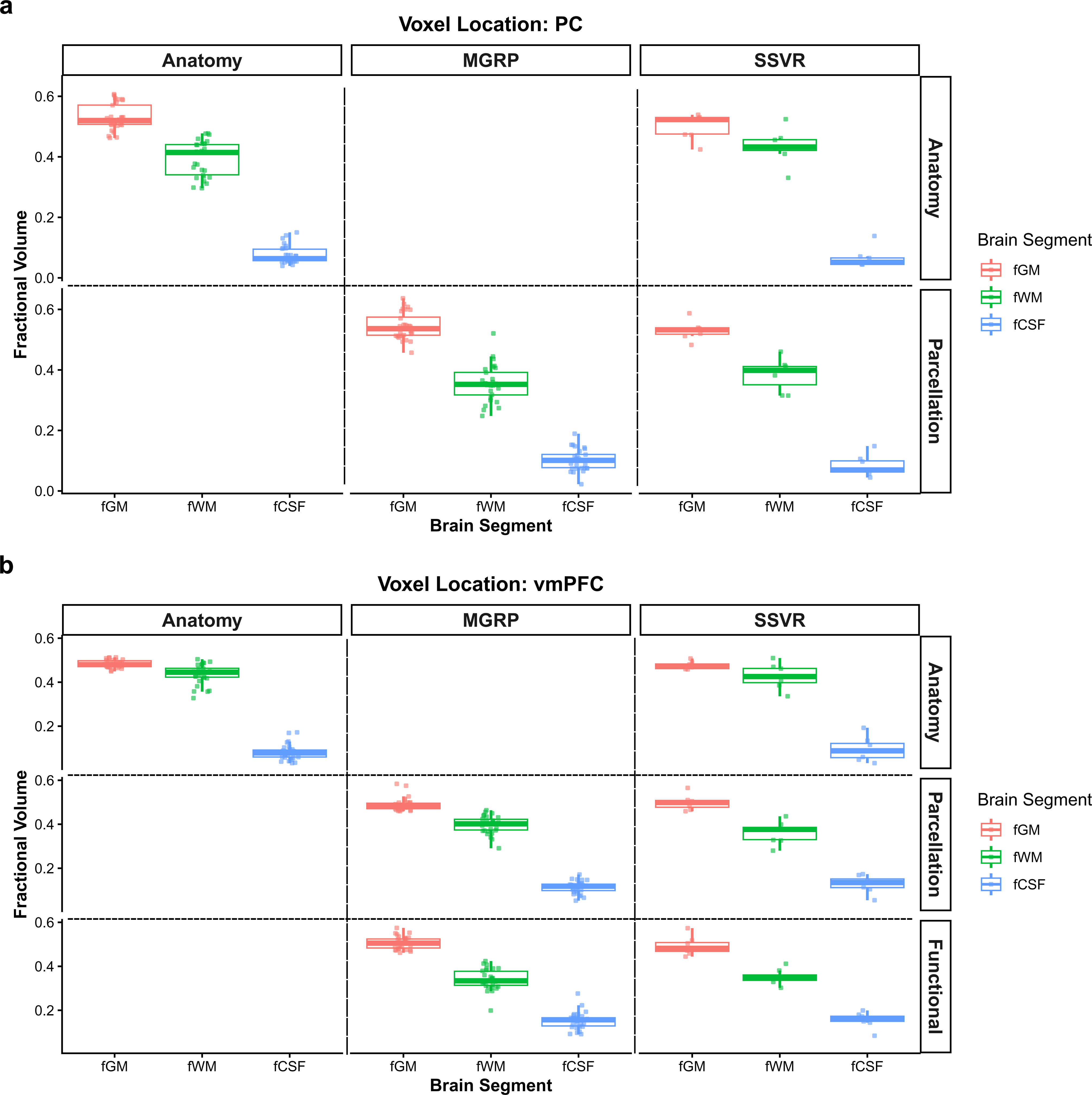
Fractional volume of the brain grey matter, white matter, and cerebrospinal fluid in the segmented MRS voxels (n=7). The figure on the top shows the fractional tissue segments at the reference benchmark regions (left parietal cortex) using the conventional anatomy-based placement, Mask-guided real-time positioning (MGRP), and Semiautomated Session-locked voxel repositioning (SSVR) methods, respectively, along with individual data points. The figure at the bottom shows fractional tissue segments at the technically challenging region (left ventromedial prefrontal cortex), also using the conventional anatomy-based placement and MGRP, and SSVR methods, respectively, along with individual data points.

Furthermore, at the technically challenging region, the ICC and within-subject CV% values improved for the SSVR method compared to the conventional anatomy-based placement and MGRP. The ICC and CV% improved for all three tissue segments (fGM, fWM, and fCSF) for all three voxels (anatomy, parcellation-mask, activation-mask). The ICC improvement was largest for the parcellation-mask voxel. For fGM ICC increased from 0.67 to 0.96, for fWM from 0.67 to 0.93, and for fCSF from 0.51 to 0.67. Furthermore, within-subject CV% decreased for all tissue segments and voxels. On average, CV% reduced from ∼3.32 to 1.45, 7.38 to 4.53 and 24.47 to 16.14 for fGM, fWM and fCSF, respectively (See to Tables 1 and S2, and Fig. 5c and 5d).

### 3.4 Metabolite concentration reliability at the Reference Benchmark region (left parietal cortex) using the conventional anatomy-based placement, MGRP, and SSVR methods

Based on the PRESS and MEGA-PRESS exclusion criteria described in sections 2.7.1 and 2.7.2, for the reference benchmark region, one participant was excluded for the conventional anatomy-based placement and SSVR method used for conventional anatomy-based placement.

tNAA: With conventional anatomy-based placement and MGRP, overall mean concentrations were comparable between anatomical (11.91±1.51) i.u. and parcellation mask placements (11.26±1.76), i.u. although variability was higher for the parcellation mask (CV% = 15.22). Using SSVR, precision improved markedly, with CV% reduced to 10.53% (anatomical) and 9.70% (parcellation mask). MDC also decreased from 2.41 to 2.34 and 3.25 to 2.11 i.u. for the anatomical and parcellation mask, respectively, indicating greater sensitivity to neurochemical changes (Table 2, S7).

**Table 2:** Metabolite concentrations (tNAA: total N-acetylaspartate, tCr: total Creatine, GABA+: Gamma-aminobutyric acid, and Glx: Glutamate + Glutamine; all measured in i.u., institutional units) and the reliability measures (CV%: percentage coefficient of variance, MDC: Minimal detectable change) for the fit at the Reference Benchmark region (left parietal cortex) using the Conventional anatomy-based placement, Mask-guided real-time positioning (MGRP), and Semiautomated Session-locked voxel repositioning (SSVR) methods.

| Voxel Placement<br>(participants) | tNAA |  | tCr |  | GABA |  | Glx |  |
| --- | --- | --- | --- | --- | --- | --- | --- | --- |
|  | CV% | MDC | CV% | MDC | CV% | MDC | CV% | MDC |
| Anatomical (6) | 10.73 | 2.34 | 5.55 | 0.68 | 11.42 | 0.56 | 21.42 | 4.34 |
| **MGRP method** |  |  |  |  |  |  |  |  |
| Parcellation mask (7) | 15.22 | 3.25 | 2.78 | 0.34 | 8.43 | 0.44 | 28.34 | 4.83 |
| **SSVR method** |  |  |  |  |  |  |  |  |
| Anatomical (6) | 10.53 | 2.41 | 4.73 | 0.55 | NA |  | 20.98 | 4.36 |
| Parcellation mask (7) | 9.70 | 2.11 | 2.04 | 0.26 |  |  | 12.95 | 2.73 |

tCr: Mean concentrations remained highly consistent across all methods and placements (ranging from 6.23 to 6.32 i.u.). However, SSVR improved precision, for both the anatomical placement and parcellation mask, where CV% decreased from 5.55 to 4.73 and 2.78 to 2.05 and MDC from 0.68 to 0.55 i.u. and 0.34 to 0.26 i.u. for anatomical placement and parcellation mask, respectively, making tCr the most stable metabolite measured (Table 2, S7).

GABA+ and Glx: GABA+ values were only available for conventional anatomy-based placement and MGRP (mean range 2.45 to 2.63 i.u.) and were not reported for SSVR, as GABA+ quantification was not performed using this approach. MGRP reduced the GABA+ CV% from 11.42% to 8.23% while MDC changed from 0.56 to 0.44. Glx exhibited the largest variability overall, with CV% reaching 28.34% under MGRP for the parcellation mask. SSVR improved its reliability for the parcellation mask, reducing CV% to 12.95% and MCD from 4.83 to 2.73 i.u. for the parcellation mask (Table 2, S7). CV% and MDC remained similar for anatomy-based placement.

### 3.5 Metabolite concentration reliability at the technically challenging region (left vmPFC) using the conventional anatomy-based placement, MGRP, and SSVR methods

Based on the PRESS and MEGA-PRESS exclusion criteria described in sections 2.7.1 and 2.7.2, for the technically challenging region, one participant was excluded for the conventional anatomy-based placement. Using the MGRP method, four participants were excluded when voxel placement was based on the parcellation mask and three participants were excluded when voxel placement was based on the functional activation mask. For the SSVR method, two participants were excluded when voxel placement was based on either the anatomical mask or the parcellation mask, and one participant was excluded when voxel placement was based on the functional activation mask.

tNAA: With conventional anatomy-based placement and MGRP, mean concentrations were highest for the parcellation mask (9.13±10.65 i.u.), though this placement also had the highest variability (CV% = 93.48). The activation mask provided the best precision for this method (CV% = 59.64). SSVR improved precision, for both anatomical and parcellation mask placement, where CV% was reduced from 93.05 to 62.37 and 93.48 to 49.53 and MDC reduced from 9.59 to 6.75 i.u. and 14.51 to 11.00 i.u., respectively. While the SSVR activation mask showed higher variability (CV% = 62.32), the anatomical and parcellation placements under SSVR generally favoured better reliability than conventional anatomy-based placement and MGRP (Table 3, S8).

**Table 3:** Metabolite concentrations (tNAA: total N-acetylaspartate, tCr: total Creatine, GABA+: Gamma-aminobutyric acid, and Glx: Glutamate + Glutamine; all measured in i.u., institutional units) and the reliability measures (CV%: percentage coefficient of variance, MDC: Minimal detectable change) for the fit at the Technically challenging region (left vmPFC) using the Conventional anatomy-based placement, Mask-guided real-time positioning (MGRP), and Semiautomated Session-locked voxel repositioning (SSVR) methods.

| Voxel Placement<br>(participants) | tNAA |  | tCr |  | GABA |  | Glx |  |
| --- | --- | --- | --- | --- | --- | --- | --- | --- |
|  | CV% | MDC | CV% | MDC | CV% | MDC | CV% | MDC |
| **Anatomy (6)** | 93.05 | 9.59 | 90.93 | 4.80 | 19.42 | 0.83 | 107.08 | 23.24 |
| **MGRP method** |  |  |  |  |  |  |  |  |
| **Parcellation mask (3)** | 93.48 | 14.51 | 101.78 | 43.25 | 24.42 | 1.05 | 140.26 | 18.94 |
| **Activation mask (4)** | 59.64 | 4.45 | 91.71 | 6.32 | 31.74 | 1.70 | 64.60 | 7.32 |
| **SSVR method** |  |  |  |  |  |  |  |  |
| **Anatomy (5)** | 62.37 | 6.75 | 84.92 | 10.11 | NA |  | 94.84 | 12.99 |
| **Parcellation mask (5)** | 49.53 | 11.00 | 63.23 | 27.01 |  |  | 112.62 | 21.03 |
| **Activation mask (6)** | 62.32 | 8.07 | 64.75 | 4.42 |  |  | 111.65 | 20.90 |

tCr: Using conventional anatomy-based placement and MGRP, mean concentrations varied considerably, particularly for the parcellation mask (17.59±36.29 i.u.), which exhibited extreme variability (CV% = 101.78). However, the SSVR method significantly stabilized these measurements across all placements. Precision for the parcellation mask improved, with CV% dropping from 101.78 to 63.23 and MDC decreasing from 43.25 to 27.01 i.u. The functional mask placement under SSVR yielded the most stable result for this metabolite (CV% = 64.75, MDC = 4.42) (Table 3, S8).

GABA+: GABA+ values were only available for the conventional anatomy-based placement and MGRP method and were not reported for SSVR. Within this method, anatomical placement showed the most stable results (CV%= 19.42 and MDC= 0.83 i.u.). In contrast, activation mask placement showed substantially higher variability (CV% = 31.74 and MDC = 1.70 i.u.) (Table 2, S7).

Glx: Glx displayed high variability overall, particularly with conventional anatomy-based placement and MGRP, where CV% reached 140.26 for the parcellation mask and 107.08 for anatomical placement. Precision improved when using the activation mask as indicated by the CV% of 64.60 (MGRP). SSVR improved the CV for anatomy (94.84%) and parcellation mask (112.62%) but not for activation mask (111.65%). Although SSVR enhanced the stability of anatomical and parcellation placements compared to conventional anatomy-based placement and MGRP, Glx remained one of the most variable metabolites, with MDC values ranging from 23.24 to 7.32 i.u (Table 2, S7).

## 4 Discussion

### Voxel placement: Global Overlap, Euclidean Distance, and Brain Segmentation

In the present study, SSVR substantially improved mean within-subject global voxel overlap, increasing from 0.64 to 0.85 at the reference benchmark region and reaching up to 0.88 at the technically challenging region across anatomical, parcellation, and activation masks compared to both conventional anatomy-based placement and MGRP methods. This suggests more consistent voxel placement and improved anatomical targeting across sessions. Geometric within-subject reproducibility was also near perfect, with within-subject Euclidean distance ICC reaching 0.99, indicating highly stable voxel centring compared to conventional anatomy-based placement and MGRP.

SSVR further improved tissue composition stability at the technically challenging region, with marked increases in ICC across fGM, fWM, and fCSF. The largest gain was observed for fGM in the parcellation-mask voxel (0.68 to 0.96), highlighting improved handling of cortical boundary regions and a more stable GM/WM balance, which is critical for longitudinal MRS studies. Despite these improvements in within-subject reproducibility, CV% remained largely similar across fGM and fWM methods for the reference benchmark regions but reduced greatly with SSVR for the technically challenging region, suggesting that residual variability in the reference benchmark region is more likely driven by physiological fluctuations or region-specific technical factors as localized B_0_ magnetic field drift, subtle shimming instabilities, and RF coil sensitivity variations rather than voxel placement.

Across the literature, a consistent hierarchy of voxel placement reproducibility emerged, progressing from manual conventional anatomy-based placement and MGRP approaches to increasingly automated solutions. Manual placement methods demonstrated only moderate spatial reproducibility, with reported within-subject overlap around 86%±5% and between-subject overlap of approximately 75%±10%, reflecting sensitivity to anatomical variability and differences in voxel precision between operators despite controlled acquisition conditions (Bai et al., 2017). Similarly, manual voxel placement at 7T showed significantly lower reproducibility compared to automated approaches, with geometric overlap ratios of ∼0.70 and significant voxel displacement across sessions (Dou et al., 2015).

In contrast, automated and semiautomated pipelines substantially improved spatial consistency, with voxel overlap increasing to ∼0.91 and reduced displacement across repeated scans, alongside improved reproducibility over time. More advanced fMRI-guided automated placement frameworks further extended these improvements by demonstrating consistently higher voxel overlap, reduced CoG displacement, and lower variability in tissue composition across both short-term and longitudinal acquisitions, while maintaining stable mean GM/WM fractions but reducing their variance (Bishop et al., 2023). However, these approaches still exhibited limitations, including residual manual intervention for cortical adjustment, restricted validation primarily in the left DLPFC, and dependence on accurate registration pipelines. Within this context automated voxel placement (Woodcock et al., 2018) demonstrated very high within-subject reproducibility (∼97% overlap) and strong between-subject consistency (>94%), but remained sensitive to small co-registration errors due to dependence on a brain template that could affect tissue composition, particularly in gray matter fractions.

Extending beyond these methods, our SSVR shows comparable or improved performance across multiple anatomical and mask-based conditions, with high voxel overlap (up to ∼0.88), near-perfect geometric within-subject reproducibility (ICC ∼0.99), and improved stability of tissue segmentation (e.g., fGM ICC up to 0.96), suggesting enhanced robustness to anatomical boundary effects and improved consistency of voxel definition across heterogeneous conditions and making the voxel overlap comparable to that of well-defined anatomical regions (∼87% to 90%) (Hehl et al., 2025). Overall, the literature supports a clear progression in methodological robustness, from conventional anatomy-based placement and MGRP to automated voxel placement and more recent automated frameworks, culminating in approaches like SSVR that use prior knowledge of the voxel placement of the original scan, resulting in improved geometric within-subject reproducibility and tissue-level stability, thereby improving reproducibility for longitudinal and multi-region MRS applications.

### Metabolite concentration measurement

In the present study, the reference benchmark region showed high baseline metabolite stability, with CV% values for tNAA and tCr consistently within 10–15%, indicating robustness to placement variability. SSVR improved between-session reproducibility across both regions. In the reference benchmark region, it refined already stable measurements, reducing tNAA CV from ∼15% to ∼10% (∼33% improvement). In contrast, the technically challenging region exhibited substantially higher variability, with CV values for tCr and Glx exceeding 170–200% under conventional anatomy-based placement and MGRP, likely due to susceptibility effects from nearby sinuses. In addition, increased variability in spectral fitting, particularly related to baseline estimation in the presence of imperfect water suppression and baseline distortions, may have contributed to the observed variability, even when the underlying spectra were relatively similar. These findings suggest that while manual placement may be sufficient in stable regions like the reference benchmark region, the observed improvement in CV achieved with the SSVR method at this region indicates that more advanced registration approaches can still enhance placement consistency. Furthermore, anatomically complex regions, such as the technically challenging region, are likely to benefit even more from the advanced registration methods. The SSVR method produced substantial improvements in between-session reproducibility, with reductions in CV of approximately 70% for tCr, 59% for tNAA, and 51% for Glx. These improvements are expected because, unlike conventional anatomy-based placement and MGRP, SSVR incorporates voxel placement information from the original scan, enabling more consistent reproduction of the same voxel location across repeated examinations. This indicates that SSVR minimizes voxel misplacement into high-noise or CSF-rich regions, which is particularly important in inhomogeneous areas.

Comparison of different MRS regions highlights a clear regional dependence of MRS between-session reproducibility, consistent with the present findings. In the literature, relatively stable regions such as the anterior cingulate cortex (ACC) and motor cortex show low variability (CV ∼5–12%) across multiple metabolites, including Glx, tNAA, tCr (Bell et al., 2021; Wang et al., 2024), while the posterior cingulate cortex demonstrates similarly low values (CV = 2.9% for Glx and 8.8% for GABA) (Baeshen et al., 2020). Motor cortex measurements also remain stable (∼6–10%), even for edited metabolites such as GABA (Gajdošík et al., 2021; Zhang et al., 2018). These findings align with the low variability observed at the PC in the current study. CV% achieved with the conventional anatomy-based placement and MGRP methods improved with SSVR for tNAA, tCr, and Glx (∼12–15% for tNAA lowering to ∼10% with SSVR, ∼4-9% for tCr lowering to ∼2-6%), suggesting that even in robust regions, SSVR can further enhance between-session reproducibility.

In contrast, more complex or susceptibility-prone regions, such as the hippocampus and frontal cortex, exhibit higher variability, with CV values of ∼10–25% for Glx and up to ∼20–50% or more for other metabolites in clinical cohorts (Gajdošík et al., 2021; Mikkelsen et al., 2019; Vigneswaran et al., 2015; Yasen et al., 2017). In comparison, the vmPFC results in the present study, particularly under conventional anatomy-based placement and MGRP, where CV values exceeded 101% for tCr, and 140% for Glx represent an extreme case. This likely reflects the combined effects of susceptibility artifacts, voxel misplacement, exclusion of multiple participants due to low data quality and likely that the sessions were 2-3 weeks apart which could have led to differences in resting-state of the participants. Importantly, SSVR substantially reduced this variability, lowering CV values to ∼63–85% for tCr and ∼49% for tNAA, bringing them closer to the upper range reported in challenging regions. While SSVR cannot fully overcome intrinsic limitations such as magnetic field inhomogeneity or differences in the resting state activity, it effectively mitigates placement-related variability, a major contributor to the extreme values observed under conventional anatomy-based placement and MGRP. The continued high variability of Glx, both in the literature and in the present study, further highlights the impact of spectral complexity and low signal-to-noise ratio, rather than systematic physiological fluctuations, given that low variability was concurrently observed for the same data within the reference benchmark region.

Beyond CV-based estimates of between-session reproducibility, MDC provides an additional measure of absolute reliability by quantifying the smallest individual change that can be interpreted as exceeding measurement error (Haley and Fragala-Pinkham, 2006; Weir, 2005). However, MDC remains rarely reported in MRS reproducibility studies, where CV, ICC, and Bland–Altman analyses are more commonly used to characterize measurement stability. Consequently, direct comparisons of MDC values across MRS studies remain limited. Riemann et al. established an MDC framework for single-voxel proton MRS and reported MDC values ranging approximately from 0.40 to 2.23 μmol/g depending on metabolite and acquisition conditions, demonstrating that detectable metabolite changes are strongly influenced by acquisition strategy and session-to-session variability (Riemann et al., 2022). Although the absolute values cannot be directly compared due to differences in units, acquisition protocols, and quantification approaches, these findings provide an important reference for interpreting measurement error in MRS. Consistent with the reduction of CV, SSVR decreased MDC values for tNAA for both, conventional anatomy-based placement, and the MGRP method (anatomy: 9.59 to 6.75 i.u.; MGRP: 14.51 to 11.00 i.u.), and tCr for the MGRP method (43.25 to 27.01 i.u.), indicating reduced measurement error and improved sensitivity for detecting true longitudinal changes. The larger MDC values observed in the technically challenging region likely reflect the additional contribution of susceptibility effects, voxel placement variability, and spectral fitting uncertainty, which are less prominent in conventional MRS regions. Similarly, SSVR reduced Glx MDC values (overall range: 23.24 to 7.32 i.u.), although Glx remained the least stable metabolite, likely due to its greater spectral complexity, lower signal-to-noise ratio, and increased fitting uncertainty.

Furthermore, for edited GABA measurements, previous reliability studies have generally reported CV and ICC rather than MDC values, limiting direct comparison of absolute measurement error. Larger variability has been reported in the literature for edited GABA as compared to unedited sequences. For example, GABA CV values of ∼4% in the dorsolateral prefrontal cortex (DLPFC) and ∼8% in the rostral ACC have been reported (Duda et al., 2021). However, in medial parietal regions, GABA variability ranges from 16.9% to 28.8% (Mikkelsen et al., 2019) and ∼12–27.6% (Mikkelsen et al., 2016). In the DLPFC, CV values as high as 27.9% have been observed (Yasen et al., 2017), while ACC measurements range between 12.6% and 14.8% (Yasen et al., 2017). In the current study, MDC analysis for GABA+ was limited to conventional anatomy-based placement and MGRP, where anatomical placement demonstrated lower MDC than activation-based placement (0.83 vs. 1.70 i.u.), consistent with the corresponding CV values. These findings suggest that the voxel placement strategy influences not only relative reproducibility but also the absolute error associated with edited metabolite measurements.

Together, these findings highlight that even in commonly studied cortical regions, GABA between-session reproducibility can span from low (∼3–8%) to moderate (∼10–30%), depending on acquisition method, metabolite, and anatomical complexity. In line with this, the present study showed GABA CV values of ∼8–11% at the PC and substantially higher variability at the vmPFC (∼19–32%). However, as SSVR was not collected for the GABA measurements, its potential impact on reducing GABA between-session variability in these regions remains unclear.

Overall, these findings are consistent with the broader MRS literature, confirming that between-session reproducibility is strongly region- and method-dependent. The parietal cortex behaves similarly to other stable regions, whereas the vmPFC represents a more extreme and challenging environment. Crucially, the substantial reduction in variability achieved with SSVR underscores the importance of incorporating prior knowledge from previous scans for improving measurement within-subject and between-session reproducibility and absolute reliability in regions where conventional anatomy-based placement/MGRP methods are insufficient. This is particularly beneficial in regions with pronounced susceptibility gradients, where small differences in voxel position or orientation can substantially alter spectral quality and metabolite quantification. By more consistently reproducing the original voxel placement, SSVR reduces these sources of variability and improves measurement reliability by reducing within-subject error and improving voxel placement consistency.

Importantly, the roles of MGRP and SSVR are complementary rather than mutually exclusive. Both methods improved compared to conventional anatomy-based placement by incorporating subject-specific prior information, but at different stages of the workflow. MGRP improves the initial voxel definition by using individualized anatomical or functional masks, thereby enhancing both anatomical targeting and longitudinal consistency compared with conventional anatomy-based placement. SSVR further improves within-subject reproducibility by registering the follow-up scan to the original acquisition, enabling more accurate reproduction of the previously defined voxel location. Consequently, SSVR can be combined with either conventional anatomy-based placement or MGRP, allowing longitudinal studies to benefit from both improved initial voxel targeting and improved session-to-session voxel reproducibility. In contrast, for cross-sectional studies where participants are scanned only once, methods such as MGRP remain valuable for achieving a consistent voxel definition across the study cohort, whereas the principal advantage of SSVR lies in repeated measurements of the same individual.

### Strengths and limitations

A key strength of this study is its comprehensive and systematic design, which included multiple MRS acquisitions across two spectroscopic sequences (PRESS and MEGA-PRESS), two anatomically and methodologically distinct brain regions (a reference benchmark region and a technically challenging region), and multiple voxel placement strategies. This multi-dimensional framework allowed direct comparison of reliability across both “benchmark” and “technical challenge” target regions, as well as across conventional and advanced voxel localization approaches. In addition, the inclusion of metabolite-specific measures (Glx and GABA+, alongside tNAA and tCr) enabled a more robust evaluation of method performance across different spectral editing and quantification conditions. A further strength of the proposed MGRP and SSVR pipelines is their rapid online processing time, requiring approximately 5 minutes once the MPRAGE scan is available. This enables processing to be initiated immediately after anatomical acquisition, such that the resulting outputs can be readily available by the time MRS planning begins. The pipelines were implemented on a standard desktop PC equipped with 16 GB RAM and an 8-core CPU running Windows 10, with FSL executed through a virtual Linux environment. This rapid processing and straightforward computational setup support the practical feasibility of the approach and facilitate seamless integration into routine scanning workflows, including time-constrained clinical and research settings. A further strength of the proposed framework is its flexibility in implementation. Although MGRP ideally relies on pre-generated masks derived from prior structural parcellation or functional ROI analyses, and therefore benefits from an established preprocessing pipeline, this is not a strict requirement for study initiation. In cases where such resources are not available, conventional anatomy-based placement can be used for the first session, followed by the application of SSVR to ensure improved voxel placement consistency and reliability across subsequent sessions.

However, a few limitations should be considered when interpreting the findings. First, all MRS voxel placements were performed by a single operator; therefore, inter-rater reliability for the operators could not be assessed. Evaluating multiple operators would have provided insight into whether the proposed methods (MGRP and SSVR) improve consistency not only within but also across operators. Second, a potential limitation of the MGRP approach is that it relies on pre-generated masks derived from prior structural parcellation or functional ROI analyses to guide initial voxel placement. As a result, an established preprocessing pipeline is required before the first MRS session, which may slightly extend preparation time and may not yet be readily available to all research groups. However, to facilitate widespread implementation and ensure this approach is readily accessible to all research groups, we have made our complete preprocessing and placement scripts openly available (OSF: https://osf.io/7ejg3/overview). Third, the SSVR method was applied only to PRESS acquisitions and was not evaluated for MEGA-PRESS. Therefore, its performance and applicability to edited spectroscopy sequences (e.g., GABA measurements) or to other localization methods such as sLASER remain to be established. Fourth, the study was conducted exclusively in healthy participants, which may limit generalizability to clinical populations as well as elderly populations with possible larger anatomical and functional variability. Future studies should therefore evaluate these voxel placement approaches in clinical cohorts to determine their robustness and potential utility in patient populations where accurate and reproducible MRS voxel positioning is particularly challenging.

## Conclusion

Overall, both MGRP and SSVR improved voxel placement and metabolite measurement reproducibility compared with conventional anatomy-based placement by incorporating subject-specific prior information. While MGRP improved the initial voxel definition using individualized anatomical or functional masks, SSVR further enhanced within-subject between-session reproducibility by reproducing the original voxel placement across repeated examinations. Although improvements in voxel placement and metabolite quantification at the reference benchmark region were modest, both methods demonstrated advantages over conventional anatomy-based placement, with SSVR providing the greatest improvements at the technically challenging region. These reductions in variability and measurement error enhance statistical power in multisession, multicentre, and longitudinal studies by lowering measurement noise and enabling more reliable detection of biological changes, particularly in regions prone to high variability such as the vmPFC, hippocampus, and brainstem. As SSVR can be combined with either conventional anatomy-based placement or MGRP, longitudinal studies can benefit from both improved initial voxel targeting and enhanced reproducibility of voxel placement across repeated sessions.

## Data Availability

To ensure facilitation and widespread use of this approach the scripts will be made accessible to all research groups through OSF upon acceptance of the manuscript. However, the data used for the study will be made available on request.

## CRediT authorship contribution statement

**Harleen Chhabra:** Writing – review & editing, Writing – original draft, Visualization, Investigation, Formal analysis, Data curation, Conceptualization, Funding. **Melina Hehl:** Writing – review & editing, Visualization. **Koen Cuypers:** Writing – review & editing, Visualization. **Ulrike Dydak:** Writing – review & editing, Visualization. **Michael A. Nitsche:** Writing – review & editing, Supervision, Project administration, Methodology, Funding acquisition. **Erhan Genç:** Writing – review & editing, Supervision, Methodology, Conceptualization. **Michael Burke:** Writing – review & editing, Visualization, Methodology, Conceptualization.

