## Supplementary material for "Testing the reliability of novel Voxel Placement approaches for Magnetic Resonance Spectroscopy": Supplemetary Document

**Supplementary Document 1**

**MRSinMRS Checklist**

| 1. Hardware | | | |
| --- | --- | --- | --- |
| a. Field strength (T) | 3 T | | |
| b. Manufacturer | Siemens | | |
| c. Model | ve syngo MR E11 | | |
| d. RF coils | 64 channels ^1^H head coil | | |
| 2. Acquisition | | | |
| a. Pulse sequence | Glx: PRESS | | GABA: MEGA-PRESS |
| b. VOI locations | Left parietal cortex and left ventromedial prefrontal cortex | | |
| c. VOI size (mm^3^) | 2x1.5x3 cm^3^ | | |
| d. TR/TE (ms) | Glx: 2000/30 | | 2000/68 |
| e. Averages | Glx: 225 | | GABA: 128 (64 OFF / 64 ON) |
| f. spectral width in Hz, duration, editing pulse 1 and 2 in ppm, editing pulse width in Hz | Glx: 2000, 1024, 0, 0 | | GABA: 2000, 1024, 7.5 and 1.9, 5.8 |
| g. Water suppression method | VAPOR | | |
| h. Shimming method | FastestMap | | |
| 3. Data analysis methods and outputs | | | |
| a. Analysis software | Glx: Osprey 2.8.1 | | GABA: Gannet 3.5.1 |
| b. Processing steps deviating from the quoted reference or product | Default Osprey with Subspectral alignment as L2Norm | | Default Gannet with eddy current correction turned on |
| c. Output measure | Raw tNAA, tCr, Glx | | GABA+ |
| d. Quantification references and assumptions, fitting model assumptions | Basis set includes default 19 simulated metabolites + measured macromolecule. The baseline knot spacing was set to 0.5 ppm, and the data were fitted with Osprey fitting | |  |
| 4. Data quality | | | |
| a. SNR (Cr), linewidth (Cr) [Hz] | Across two voxel localizations and placement methods. Refer to Table S1-4 for details | | |
| b. Data exclusion criteria | CrSNR and/or CrFWHM ≤ 2±SD, CRLB < 20% | GABA+ SNR and/or GABA+ FWHM ≤ ±2SD, GABA Error Fit < 20% | |
| c. Quality measures of postprocessing model fitting | CRLB | | ErrorFit |
| d. Sample spectrum | 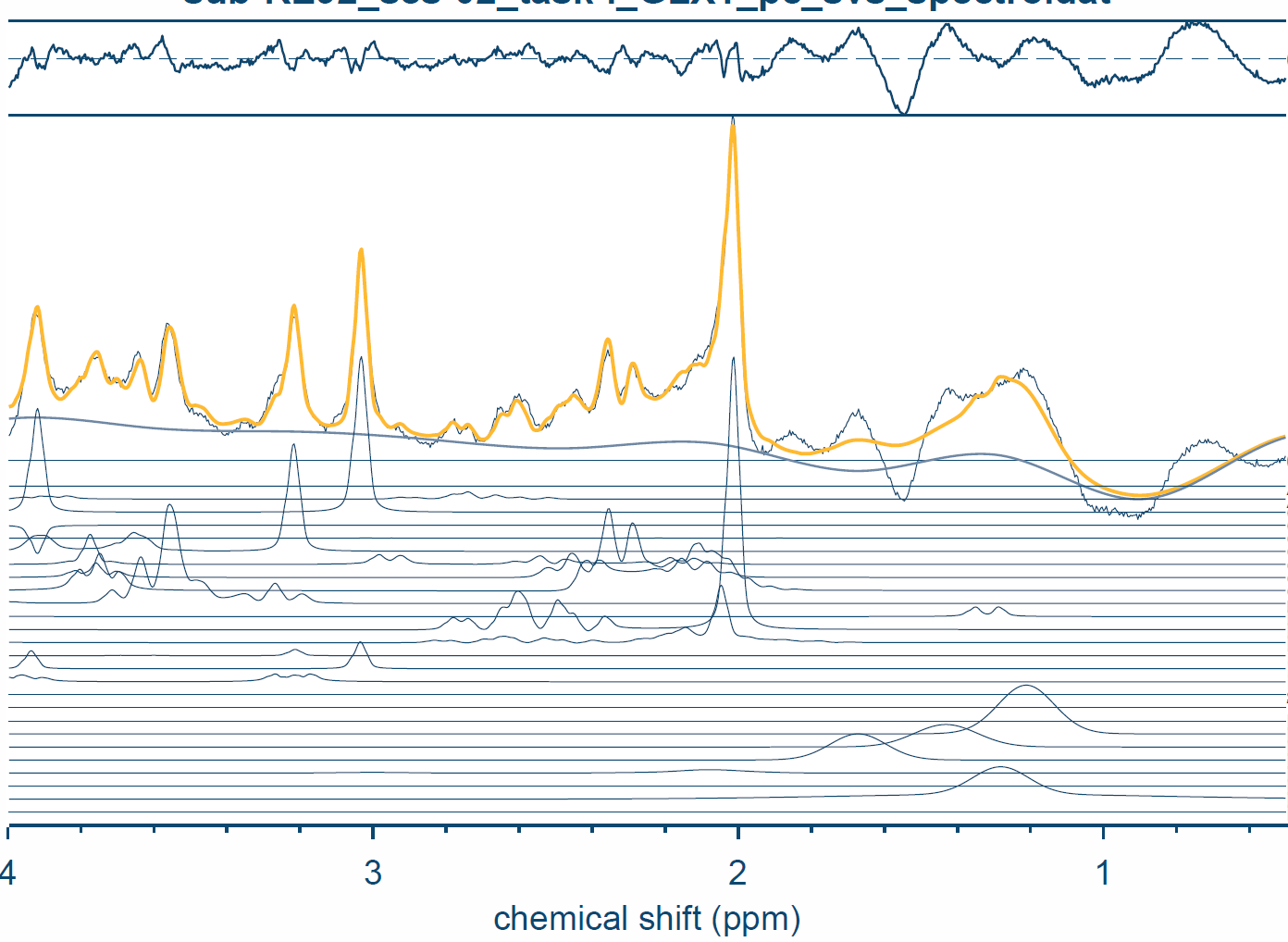 | | 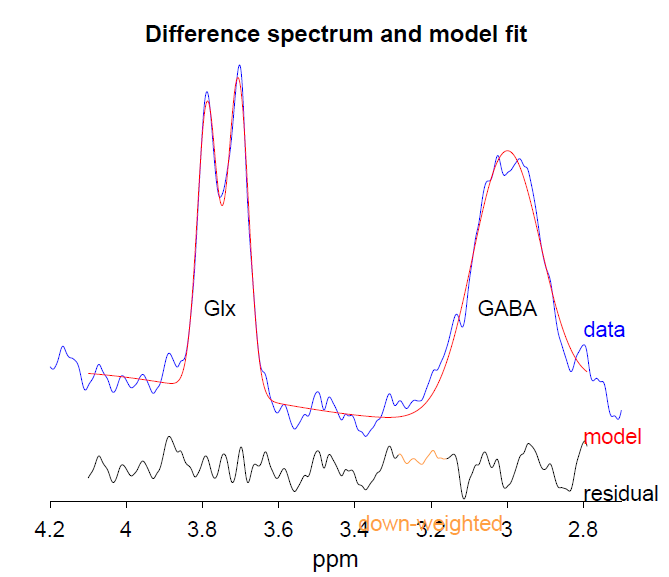 |

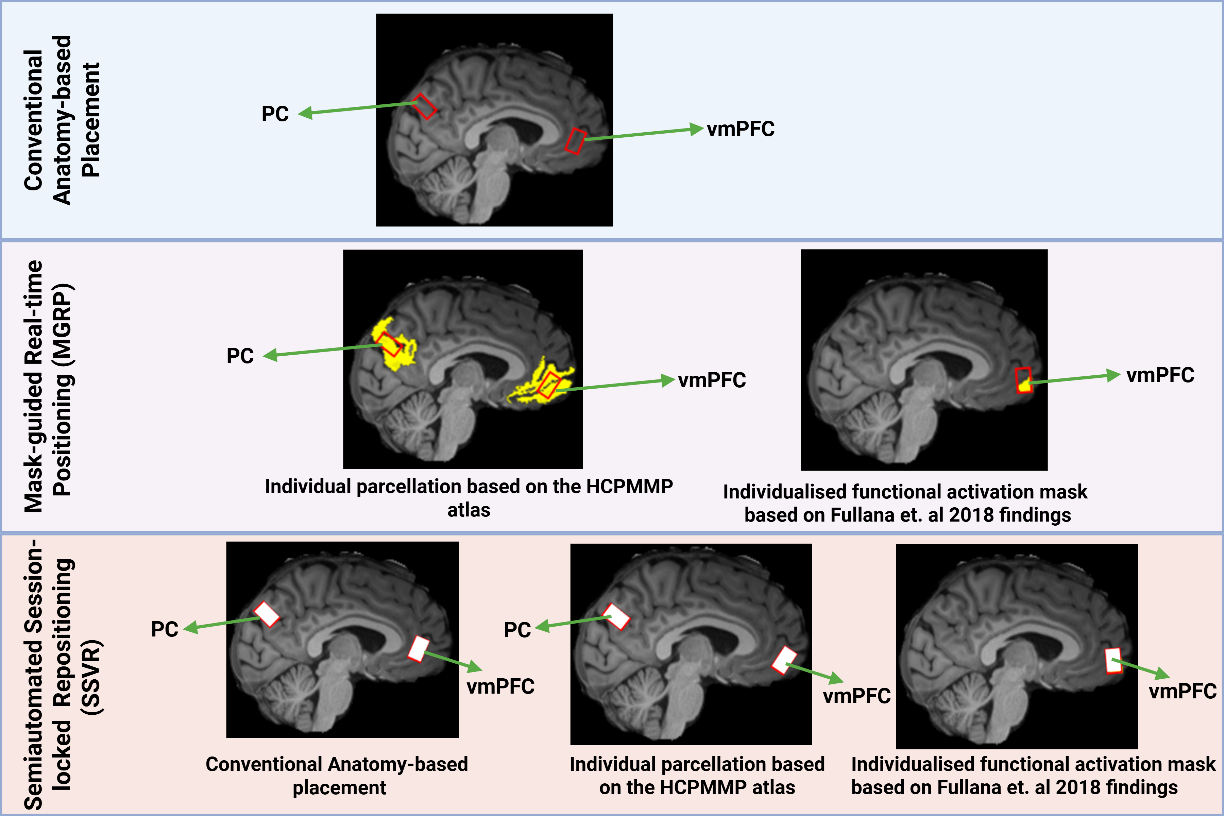

**Fig S1: Voxel definitions for the reference benchmark region, the left parietal cortex (PC), and the technical challenge region, the left ventromedial prefrontal cortex (vmPFC).** Voxels for both regions were placed based on the conventional anatomical-landmark-based placement (top), individualised regional masks (middle), and previous MRS session (bottom) voxel positions. For both PC and vmPFC, an individualised anatomical mask was created with the HCPMMP atlas parcellation in Freesurfer. For the vmPFC, an additional individualized functional mask was created by linearly transforming a task-based activation mask derived from the study by Fullana et al. (2018) from MNI space into each participant's native anatomical space.

Table S1: The overall fractional tissue mean and standard deviation values of the voxel at the reference benchmark region (parietal cortex) using the Conventional anatomy-based placement, Mask-guided real-time positioning (MGRP), and Semiautomated Session-locked voxel repositioning (SSVR) methods.

| **Voxel placement (participants)** | **fGM** | **fWM** | **fCSF** |
| --- | --- | --- | --- |
|  | **Mean±SD** | **Mean±SD** | **Mean±SD** |
| **Anatomy (7)** | 0.53±0.04 | 0.39±0.06 | 0.08±0.03 |
| **MGRP methods** | | | |
| **Parcellation mask (7)** | 0.54±0.04 | 0.35±0.04 | 0.10±0.04 |
| **SSVR method** | | | |
| **Anatomy (7)** | 0.52±0.05 | 0.41±0.06 | 0.07±0.03 |
| **Parcellation mask (7)** | 0.53±0.04 | 0.38±0.06 | 0.09±0.04 |

Table S2: The overall fractional tissue Mean and standard deviation values of the voxel at the reference benchmark region (parietal cortex) using the Conventional anatomy-based placement, Mask-guided real-time positioning (MGRP), and Semiautomated Session-locked voxel repositioning (SSVR) methods.

| **Voxel placement (participants)** | **fGM** | **fWM** | **fCSF** |
| --- | --- | --- | --- |
|  | **Mean±SD** | **Mean±SD** | **Mean±SD** |
| **Anatomy (7)** | 0.48±0.02 | 0.44±0.05 | 0.08±0.04 |
| **MGRP methods** | | | |
| **Parcellation mask (7)** | 0.49±0.03 | 0.40±0.04 | 0.11±0.03 |
| **Activation mask (7)** | 0.51±0.03 | 0.34±0.05 | 0.15±0.04 |
| **SSVR method** | | | |
| **Anatomy (7)** | 0.48±0.02 | 0.43±0.05 | 0.09±0.05 |
| **Parcellation mask (7)** | 0.50±0.04 | 0.37±0.06 | 0.12±0.04 |
| **Activation mask (7)** | 0.50±0.04 | 0.35±0.04 | 0.15±0.04 |

Table S3: Quality parameters for MEGA-PRESS GABA+ fit at the reference benchmark region (parietal cortex) using the Conventional anatomy-based placement, Mask-guided real-time positioning (MGRP), and Semiautomated Session-locked voxel repositioning (SSVR) methods.

| **Voxel Placement (participants)** | **GABA+ error fit** | **GABA+ SNR** | **GABA+ FWHM** |
| --- | --- | --- | --- |
|  | **Mean±SD** | **Mean±SD** | **Mean±SD** |
| **Anatomical (7)** | 5.68±1.19 | 9.37±1.16 | 22.3±2.19 |
| **MGRP methods** | | | |
| **Parcellation mask (7)** | 5.82±1.39 | 9.12±0.78 | 23.35±1.85 |
| **SSVR method** | | | |
| NA | | | |

Table S4: Quality parameters for PRESS Glx fit at the reference benchmark region (parietal cortex) using the Conventional anatomy-based placement, Mask-guided real-time positioning (MGRP), and Semiautomated Session-locked voxel repositioning (SSVR) methods.

| **Voxel Placement (participants)** | **Cr SNR** | **Cr FWHM** | **Water FWHM** | **Glx CRLB** |
| --- | --- | --- | --- | --- |
|  | **Mean±SD** | **Mean±SD** | **Mean±SD** | **Mean±SD** |
| **Anatomy (6)** | 74.22±52.13 | 5.65±0.35 | 6.43±0.24 | 11.58±9.45 |
| **MGRP methods** | | | | |
| **Parcellation mask (7)** | 63.45±42.16 | 5.74±0.50 | 6.57±0.17 | 8.79±3.06 |
| **SSVR method** | | | | |
| **Anatomy (6)** | 67.38±47.34 | 5.78±0.66 | 6.50±0.29 | 9.33±2.91 |
| **Parcellation mask (7)** | 55.42±39.45 | 6.05±0.99 | 6.70±0.44 | 9.00±2.75 |

Table S5: Quality parameters for MEGA-PRESS GABA+ fit at the technically challenging region (ventromedial prefrontal cortex) using the Conventional anatomy-based placement, Mask-guided real-time positioning (MGRP), and Semiautomated Session-locked voxel repositioning (SSVR) methods.

| **Voxel Placement (participants)** | **GABA+ error fit** | **GABA+ SNR** | **GABA+ FWHM** |
| --- | --- | --- | --- |
|  | **Mean±SD** | **Mean±SD** | **Mean±SD** |
| **Anatomical (5)** | 7.08±2.4 | 8.55±2.75 | 28.98±8.03 |
| **MGRP method** | | | |
| **Parcellation mask (6)** | 8.33±3.79 | 7.63±1.60 | 24.38±3.23 |
| **Activation mask (2)*** | 18.48±17.44 | 39.66±15.86 | 21.57±24.53 |
| **SSVR method** | | | |
| NA | | | |

*Participants were excluded if either of the two sessions did not pass the quality check

Table S6: Quality parameters for PRESS Glx fit at the technically challenging region (ventromedial prefrontal cortex) using the Conventional anatomy-based placement, Mask-guided real-time positioning (MGRP), and Semiautomated Session-locked voxel repositioning (SSVR) methods.

| **Voxel placement (participants)** | **Cr SNR** | **Cr FWHM** | **Water FWHM** | **Glx CRLB** |
| --- | --- | --- | --- | --- |
|  | **Mean±SD** | **Mean±SD** | **Mean±SD** | **Mean±SD** |
| **Anatomy (6)** | 78.75±19.00 | 15.48±8.55 | 12.55±6.19 | 9.75±3.28 |
| **MGRP methods** | | | | |
| **Parcellation mask (3)** | 122.28±13.76 | 26.95±34.70 | 18.60±17.06 | 12.33±5.72 |
| **Activation mask (4)** | 91.03±53.11 | 14.85±5.62 | 11.81±2.58 | 9.75±4.89 |
| **SSVR method** | | | | |
| **Anatomy (7)** | 69.42±23.32 | 14.06±8.85 | 12.9±7.12 | 9.71±2.89 |
| **Parcellation mask (5)** | 57.68±18.45 | 11.97±3.92 | 10.16±0.87 | 13.00±7.86 |
| **Activation mask (6)** | 132.94±62.09 | 16.69±7.89 | 13.79±3.83 | 11.58±6.76 |

Table S7: The overall metabolite concentration mean and standard deviation values of the voxel at the reference benchmark region (parietal cortex) using the Conventional anatomy-based placement, Mask-guided real-time positioning (MGRP), and Semiautomated Session-locked voxel repositioning (SSVR) methods.

| **Voxel Placement (participants)** | **tNAA** | **tCr** | **GABA** | **Glx** |
| --- | --- | --- | --- | --- |
|  | **Mean±SD** | **Mean±SD** | **Mean±SD** | **Mean±SD** |
| **Anatomical (6)** | 11.91± 1.51 | 6.24± 0.59 | 2.45± 0.34 | 11.45± 3.53 |
| **MGPR methods** | | | | |
| **Parcellation mask (7)** | 11.26± 1.76 | 6.23± 0.26 | 2.63± 0.29 | 10.12± 4.03 |
| **SSVR method** | | | | |
| **Anatomical (6)** | 11.36± 1.14 | 6.23± 0.43 | NA | 10.74± 3.53 |
| **Parcellation mask (7)** | 11.16± 1.22 | 6.32± 0.16 |  | 11.68± 4.03 |

Table S8: The overall metabolite concentration mean and standard deviation values of the voxel at the technically challenging region (ventromedial prefrontal cortex) using the Conventional anatomy-based placement, Mask-guided real-time positioning (MGRP), and Semiautomated Session-locked voxel repositioning (SSVR) methods.

| **Voxel Placement (participants)** | **tNAA** | **tCr** | **GABA** | **Glx** |
| --- | --- | --- | --- | --- |
|  | **Mean±SD** | **Mean±SD** | **Mean±SD** | **Mean±SD** |
| **Anatomy (6)** | 6.14±4.76 | 3.48±2.79 | 2.16±0.48 | 10.20±15.49 |
| **MGPR methods** | | | | |
| **Parcellation mask (3)** | 9.13±10.65 | 17.59±36.29 | 1.99±0.73 | 6.96±12.16 |
| **Activation mask (4)** | 4.04±2.44 | 4.11±3.24 | 2.53±2.49 | 6.62±3.66 |
| **SSVR method** | | | | |
| **Anatomy (5)** | 8.37±3.99 | 3.74±2.25 | NA | 8.54±7.30 |
| **Parcellation mask (5)** | 9.38±6.07 | 3.96±3.18 |  | 7.66±7.35 |
| **Activation mask (6)** | 8.29±7.22 | 4.71±3.03 |  | 7.22±4.42 |
